# Topological constraints and finite-size effects in quantitative polymer models of chromatin organization

**DOI:** 10.1101/2023.06.16.545312

**Authors:** Amith Z. Abdulla, Maxime M. C. Tortora, Cédric Vaillant, Daniel Jost

## Abstract

Polymer physics simulations have provided a versatile framework to quantitatively explore the complex mechanisms driving chromosome organization. However, simulating whole chromosomes over biologically-relevant timescales at high resolution often constitutes a computationally-intensive task — while genes or other regions of biological interest may typically only span a small fraction of the full chromosome length. Conversely, only simulating the sub-chromosomal region of interest might provide an over-simplistic or even wrong description of the mechanism controlling the 3D organization. In this work, we characterize what should be the minimal length of chromosome to be simulated in order to correctly capture the properties of a given restricted region. In particular, since the physics of long, topologically-constrained polymers may significantly deviate from those of shorter chains, we theoretically investigate how chromosomes being a long polymer quantitatively affects the structure and dynamics of its sub-segments. We show that increasing the total polymer length impacts on the topological constraints acting on the system and thus affects the compaction and mobility of sub-chains. Depending on the entanglement properties of the system, we derive a phenomenological relation defining the minimal total length to account for to maintain a correct topological regime. We finally detail the implications of these conclusions in the case of several specific biological systems.

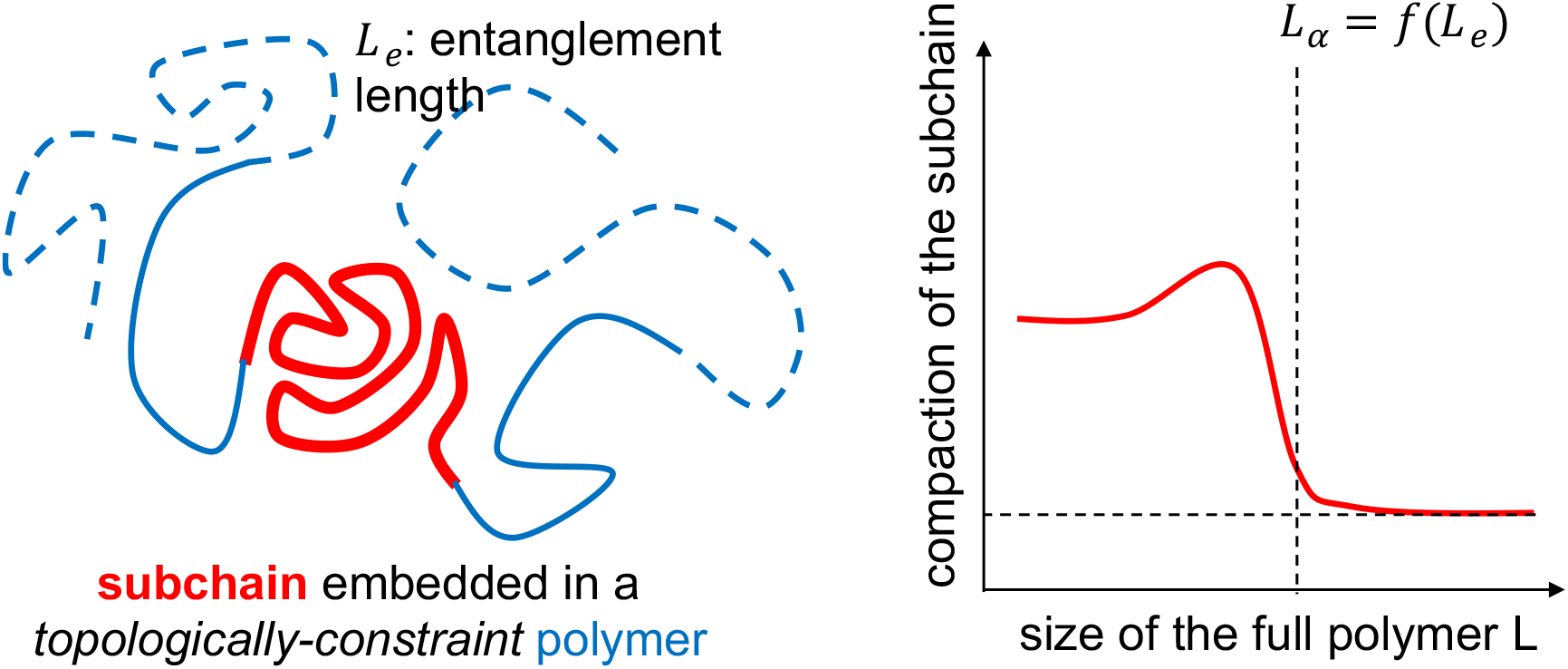

## 1 Introduction

Chromosomes are long polymers hierarchically organized at different scales during interphase [1]. At small scales (∼ 10*nm*, ∼ 200*bp*), DNA is packaged by histone proteins into the chromatin fiber. Chromatin is then organized into consecutive 3D domains with enhanced intra-domain contacts called topologically associating domains (TADs) (∼ 100s of kbp, ∼ 100*nm*). At a larger scale (∼ Mbps,∼ *μm*), genome organizes into phase-segregated, active and inactive compartments. Finally, each chromosome occupies its own territory inside the nucleus (∼ 10*μm*).

In the recent years, polymer physics has provided concrete frameworks to characterize and rationalize the mechanisms driving the spatio-temporal dynamics of chromosomes [2]: from SMC-mediated loop extrusion for TAD formation [3, 4] to epigenomic-driven interactions for chromatin 3D compartmentalization [5, 6]. With recent experimental advances, such as Micro-C [7, 8] or chromosome tracing [9, 10], it may be now possible to study specific locus, at near-nucleosome resolution (200bp-1kbp) with good statistics. Hence, more detailed polymer models are developed to finely study the organization of specific regions of interest and the interplay between the various physico-chemical mechanisms that may act locally [11].

When focusing on a region of interest, most of the models limit their considerations to a minimal chromatin domain encapsulating the corresponding genomic segment [11–13]. However, such restriction relies implicitly on the assumption that the spatial conformation of this region is not strongly affected by the rest of the chromosome(s) [17], i.e., that the folded structure of the region may be described independently of the surrounding chromatin environment. Knowing that polymer properties are sensitive to finite-size effects [14–17], a more accurate choice might be to simulate the entire chromosome on which the region resides in, which would ensure to account for all the external interactions. But, over biologically-relevant timescales, it is likely to constitute a computationally-intensive task, especially when working at high resolution. Here, we aim to theoretically investigate what would be the extent of the minimal genomic region that one should explicitly consider around a given locus in order to effectively capture correctly the dynamical and structural properties of the domain of interest and to compare with experiments.This question could be reformulated more fundamentally and generically as how chromosomes being a long polymer quantitatively affect the structure and dynamics of its sub-segments.

Typically, there may exist two major contributions of the overall polymer environment on a simulated region of interest: (i) an intrinsic contribution: the subchain (corresponding to the targeted region) is embedded inside a (often long) polymeric chain (the chromosome) that may impact the dynamical and structural properties of the subchain ; (ii) an extrinsic contribution: the subchain may have specific interactions with other regions along the long polymeric chain (or other chromosomes) that may also impact its spatio-temporal dynamics [59].

In this work, we first explore the intrinsic contribution by systematically varying the length of the polymer being simulated and quantifying the variations in structural and dynamical properties of the genomic region of interest. In particular, we show that the variations are directly linked to the topological state of the long chromosome. We can quantitatively predict the minimal length of the polymer to be simulated while studying a particular gene/locus, required to accurately capture the structure and dynamics of this region. Building on these results, we finally investigate the role of the extrinsic contribution in different biological contexts in order to devise a combined set of ground rules on how to choose the correct length to simulate.

## 2 Models and Methods

### 2.1 Lattice polymer model

We model chromatin as a semi-flexible, self-avoiding polymer using the kinetic Monte-Carlo framework developed in [17,18]. The total polymer chain is composed of *N* monomers, each of size *b* and encompassing *n* bp. The contour length of the simulated chromatin segment would be given by *L*_*c*_ = *Nb* representing *L = Nn* bp. The polymer is evolving on a face centered cubic (fcc) lattice of dimension *S* × *S*× *S* containing 4*S*^3^ lattice sites. Each lattice site can be occupied by a maximum of two monomers if and only if they are consecutive along the polymeric chain. Double occupancy of consecutive monomers accounts for the effect of contour length fluctuations [19] and allows the chain to still efficiently move in dense system. Two non-consecutive monomers cannot be at the same lattice site owing to excluded volume interactions.

Periodic boundary conditions are used to control for the volumic fraction occupied by the polymer (given by *ϕ* = *N*/4*S*^3^),and to account for the crowding effect of other chains. It is to be noted that the periodic boundary condition does not confine the polymer to the finite volume of the simulation box instead the polymer is free to extend over large distances.

The stiffness of the chain is accounted via the Hamiltonian:

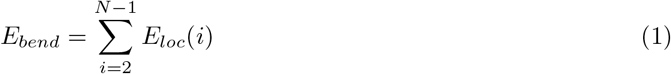

where *E*_*loc*_(*i*) = *k*_*int*_(1 *‐* cos *θ*_*i*_) [20] if there is no double occupancy around *i* and the angle *θ*_*i*_ between two successive bonds around monomer *i* is well defined (Fig.S2); otherwise, in case of double occupancy, we averaged among all the possible local bound orientations:

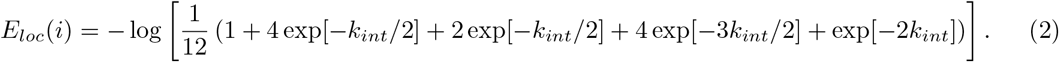

*k*_*int*_ is the bending energy and is directly related to the chain Kuhn length *L*_*k*_ [17].

To account for epigenome-driven interactions, we introduce self-attractions between monomers that share the same epigenomic state [6, 17]. This is accounted via the Hamiltonian:

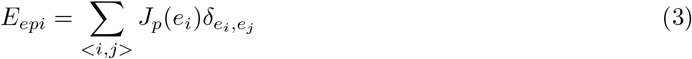

where the sum runs over all the pairs (*i, j*) of monomers occupying nearest-neighbor (NN) lattice sites, *e*_*i*_ is the epigenomic state of monomer *i*, 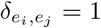 if *e*_*i*_ = *e*_*j*_ (= 0 otherwise), *J*_*p*_(*e*) is the strength of self-interaction between monomers of state *e*. All the energies are defined in units of *k*_*b*_*T* with *k*_*b*_ the Boltzmann constant and *T* the temperature.

In the following sections, unless specified otherwise (sections 2.6, 3.4), we use a fine-scale model with *n* = 1kbp, *b* = 20nm and *k*_*int*_ = 3.217k_*b*_*T* which correspond to a chromatin fiber of diameter *b*, linear compaction *≈* 50bp/nm and with a Kuhn length of *L*_*k*_ = 100nm, consistent with recent estimates in yeast [21, 22]. We maintain a chromatin volume fraction of 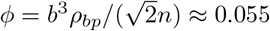, giving a bp-density of about *ρ*_*bp*_ = 0.009bp/*nm*^3^ [23] (see Table S1). With these parameters, the contour-length fluctuations arising for the double-occupancy rule are minimal (less than 2.5%, see Fig. S9) and cannot explain the observed behaviors.

### 2.2 Simulations

The dynamics of the chain is simulated using local moves on the lattice allowing the monomers to randomly hop between nearest-neighbor sites [24]. More precisely, one trial move consists in randomly picking a monomer and in attempting to move it to one of its randomly chosen nearest-neighbor, under the constraints that the chain connectivity is maintained (two consecutive monomers along the chain occupy the same or NN sites) and that the double-occupancy/self-avoidance rule (see above) is conserved (except some cases in 3.2.1). The trial move from an old configuration *o* to a new one *n* is then accepted with a probability *accept*(*o → n*) according to the Metropolis criterion on the total Hamiltonian *E*_*tot*_ *≡ E*_*bend*_ + *E*_*epi*_:

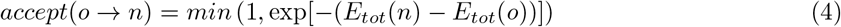

We define one Monte-Carlo step (MCS) as *N* trial monomer moves (except in section 3.2.2). Note that this local kinetic Monte-Carlo scheme prevents chain crossing and thus conserves the current topological constraints.

Numerical simulations were performed using custom made code developed in the Jost group which can be downloaded at https://github.com/physical-biology-of-chromatin/LatticePoly.

### 2.3 Initial configurations

In each situation and trajectory, we initialize the system by a “knot-free” configurations generated by the “hedgehog algorithm” [17] that do not contain knot-like entanglements. Starting from a straight chain with a few monomers, a link is randomly chosen and a monomer is inserted at site close to the two monomers such that the new configuration satisfies excluded volume and chain connectivity criterion, the process is repeated until the entire chain is grown [36]. The dynamics of the chain is then simulated during 10^8^ MCS, recording data every 10^5^ MCS. For each parameter set, we simulated at least 128 trajectories. Note that the choice of “knot-free” initial configurations is motivated by the observations that (1) chromosomes during interphase are likely to be weakly entangled and to contain very few internal knot-like structures [26–28]; (2) due to the slow large-scale reorganization kinetics of such long, topologically-constrained polymers (impossibility for two strands to cross each other), this initial state ensures that the chromosomes generally remain in a weakly-entangled/crumpled regime over the course of the simulations, as suggested by Rosa and Everaers [29].

We have also validated that the conclusions of the work do not change if we start from a “knotted” initial configuration (Fig. S4). In this case, the configuration was initialized using a random walk and thus possibly contains knot-like entanglements.

### 2.4 Strand crossing moves and disconnected chains

In section 3.2.2, to understand the effect of topology on the structure and dynamics of a region of interest, we implement strand crossing moves following the procedure described by Ubertini and Rosa [25]. One strand crossing move consists of randomly picking a monomer *i* and attempting to swap it with a random NN position if its occupied (monomer *j*) and the swap doesn’t break chain connectivity (provided *i* and *j* are not connected) (Fig.S2). On a fcc lattice, this allows for strand crossing, while 1/3 of the possible strand crossing moves are topology changing. In section 3.2.2, one MCS is redefined as *N* trial moves and *N* trial strand crossing moves (average acceptance rate is 1/1000 of the normal moves).

In section 3.2.1 (Fig. 3C,D), to explicitly study the effect of chain connectivity, we simulated at the same time two separate chains one of length *L*_*d*_ and another of length *L ‐ L*_*d*_. The two chains were created by breaking the connectivity between monomer *L*_*d*_ and *L*_*d*_ + 1 after initialization by the “hedgehog algorithm”.

### 2.5 Measured observables

For each data point, we compute several structural quantities like (1) the pairwise squared distance between monomers *i* and *j, R*^2^(*i, j*) and (2) the squared radius of gyration of a subchain [*i*_0_ : *j*_0_]:

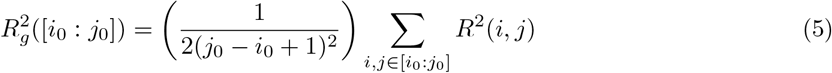

which quantifies the overall size of the region of interest (Fig. 1A). The end-to-end squared distance 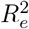 of a domain of interest *L*_*d*_ is equal to *R*^*2*^(*i*_0_, *j*_0_) with *i*_0_ and *j*_0_ the first and last monomers of *L*_*d*_, respectively.

**Figure 1.**
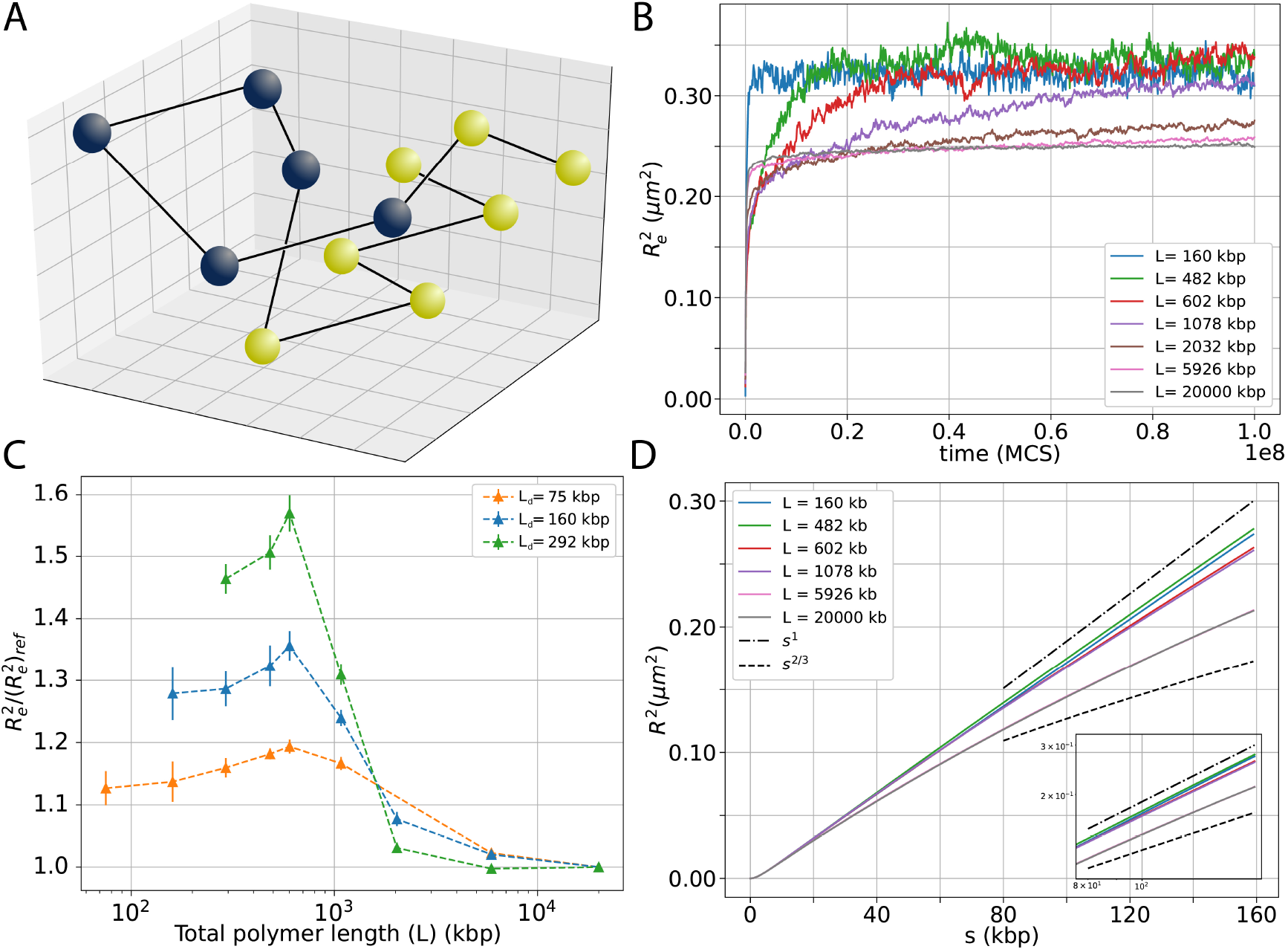
(A) An illustration of the polymer being simulated on a fcc lattice (1 monomer: 1 kbp, 20 nm). The region of interest *L*_*d*_ is schematically depicted in blue. (B) Time evolution of the squared end-to-end distance 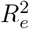 for *L*_*d*_ = 160kbp and for different total polymer lengths *L*. (C) Deviation of 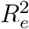 from the reference value 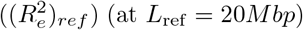 as a function of total polymer length *L* for different *L*_*d*_ values. (D) Mean squared distance *R*^2^ between monomers inside a domain of size *L*_*d*_ = 160kbp as a function of genomic distance *s* between them. Inset shows the zoom of the large genomic distances (*s* ≳ 50 kbp) in log-log scale. Note that the curves for *L* = 5926kbp and *L* = 20000kbp overlap. (B-D) In all simulations, volumic fraction ϕ = 0.055 was kept constant.

To quantify the mobility of monomers, we compute the mean squared displacement of each monomer *i*

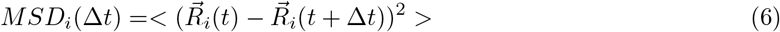

where the average is performed over time *t* and trajectories. 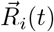 is the position of *i* in 3D at time *t*, and Δ*t* is a time-lag. The individual monomer MSD *g*_*1*_ typically corresponds to the squared distance travelled by a monomer during the time lag Δ*t* and is computed by averaging *MSD*_*i*_ over all the monomers in *L*_*d*_. We also estimate the MSD for the center of mass *g*_*3*_ of a subchain [*i*_0_ : *j*_0_]

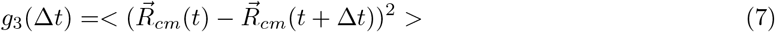

with 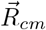 the position of the center of mass of the subchain

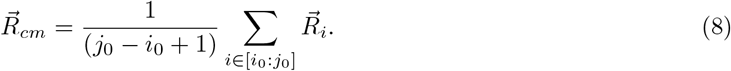

Similarly, the MSD of end-to-end vector of the domain *MSD*_*Re*_ is given by

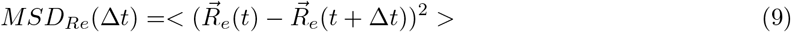

with 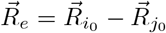 the end-to-end vector.

### 2.6 Contextualizing the simulations to specific experimental systems

We obtained processed Hi-C maps from [30] and [31] at 800-bp resolution, for Drosophila and yeast, respectively. Therefore, in section 3.4 (Fig.7), the resolution of the polymer model is set to 800-bp (*b*= 15*nm, k*_*int*_ = 4.05*k*_*b*_*T* for L_*k*_ = 100*nm*) in order to match the experimental resolution of the HiC contact data. We use a bp-density of *ρ*_*bp*_ = 0.009*bp/nm*^*3*^ for Drosophila and *ρ*_*bp*_ = 0.005*bp/nm*^3^ for yeast (see Table S1). Experimental Hi-C data were then normalized as in [32] to transform contact frequencies into contact probabilities in order to be comparable with model predictions. Then, we select one region of interest for each species: for drosophila, we consider a ∼ 307 kb region containing a topologically associating domain (TAD) [33, 34] (from 17, 955 to 18, 262 kbp) in chromosome 3R (lower part, Fig. 7), and for yeast a ∼ 50 kb-long region (from 334, 274 to 378, 470 bp) in chromosome 12 containing the gene REA1 (lower part, Fig. S6). From the experimental Hi-C maps, we identify the relevant compartments in Drosophila and domains in yeast using the “hic data.find compartments” tool from the TADbit suite [35]. In the yeast case, we assign each monomer within the domain of REA1 gene to state *e*_*M*_ — which may self-interact (*J*_*p*_(*e*_*M*_) *≡ J) —* and attribute all the remaining monomers to a non-attractive state *e*_*U*_ (*J*_*p*_(*e*_*U*_) = 0). In the Drosophila case, each monomer along chromosome 3R corresponding the B compartment is assigned a self-interacting state *e*_*M*_ (*J*_*p*_(*e*_*M*_) *≡ J*) and the other monomers a neutral state *e*_*U*_ (*J*_*p*_(*e*_*U*_) = 0). For each *J* value, we simulate 48 replicates.

To compare model predictions to experiments, we compute from the simulations the average contact probabilities between any pairs of monomers. A parameter involved in this analysis is the so-called radius of capture *R*_*c*_ that corresponds to the maximal 3D distance between two monomers captured in Hi-C experiments as a contact [36]. This is an unknown parameter, which is expected to strongly depend on the experimental protocol employed for the generation of the reference Hi-C data [32]. The structural quantities like the mean squared end-to-end distance 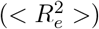 and the Hi-C maps of the regions were computed by averaging over the replicates and over the 10^7^ MCS following a relaxation step of 10^7^ MCS.

To fit the experimental Hi-C data, we systematically vary *R*_*c*_ from 15 to 150 nm and the interaction strength *J* from 0 to ‐0.4 *k*_*b*_*T*. The best-fit parameter combination (*J* and *R*_*c*_) is obtained by minimizing the *χ*^2^ score, defined as where *HiC*_*pred*_(*i, j*) is the simulated contact frequency between monomers *i* and *j HiC*_*exp*_(*i, j*) the corresponding experimental data, and *t* is the ensemble of pairs of monomers on which the fit is based, with cardinal number *N*_*t*_. In our case, *t* is limited to all pairs of monomers with a genomic distance larger than 4kbp, as our coarse-grain model is not expected to correctly capture local, fine-scale details below 4kbp. The corresponding parameter values are verified to also maximize the Pearson correlation between the experimental and simulated Hi-C matrix (Fig. S7,S8), thus evidencing the accuracy of our optimization protocol.

To quantify the degree of compartmentalization in the region of interest, we also compute the saddle-strength compartment score values *S* using cooltools (https://cooltools.readthedocs.io/en/latest/notebooks/compartments_and_saddles.html#Saddle-strength)), which is related to the ratio of the contacts within same compartments by the contacts between different compartments.

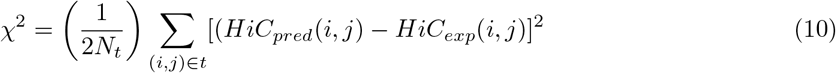

## 3 Results

### 3.1 Total polymer length determines domain compaction and mobility

We start by investigating the intrinsic contribution of polymer environment in the simple case of a homopolymer purely driven by self-avoidance, starting from a compact, post-mitotic-like, knot-free configuration.

#### 3.1.1 Impact on compaction

We consider a domain of interest *d* of size *L*_*d*_ embedded within a larger polymer of total size *L* (Fig. 1A). To specifically understand the effects of changing length, for a given domain size *L*_*d*_, we simulate different systems by varying *L* (from *L* = *L*_*d*_ to *L* = 20*Mbp*, the typical size of a chromosome arm in Drosophila [37]) keeping the volume fraction *ϕ* constant to conserve volumic density (see Table S1). We consider the simulations performed for *L*_*ref*_ = 20Mbp as the reference case and investigate whether the structural and dynamical reference properties of *d* are conserved when *L* is decreased. For simplicity, we place the domain of interest in the middle of the larger polymer but this position does not impact the results, unless it is placed at the very end of the polymer (Fig. S1).

As a first step, we study how *L* affects the structural characteristics for *L*_*d*_ = 160kbp, a size much smaller than the length of our reference case (*L*_ref_ = 20Mbp). We follow the dynamics of decompaction of the initial configuration and of convergence towards a (pseudo)stationary state for structural properties of the domain d. In particular, we measure the ensemble-averaged end-to-end squared distance (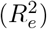) of d (see 2.5) as a function of time (Fig. 1B). Such an observable corresponds to the slowest Rouse mode of a polymer chain [38] and thus well captures the full convergence towards equilibrium. We also compute the time evolution of the squared radius of gyration (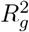) (Fig. S3). For small *L* (< 500kbp), we observe a rapid expansion of the domain and a fast convergence (less than 10^7^ MCS) towards an equilibrium state. For large *L* (> 2*Mbp*), there is still a first rapid decompaction step but followed by a slower convergence (∼ 5 *×* 10^7^ MCS) towards a pseudo-stationary state with 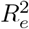 being almost constant but still very slowly and slightly expanding at long time-scale (> 10^8^ MCS). Between these two regimes (*L* = 1.078Mbp in Fig. 1B) the system displays a very slow convergence towards the steady state. Interestingly, for *L* ≲ 1 Mbp all the curves converge towards approximately the same value 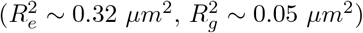, while, for *L* ≳ 2Mbp, they converge towards a more compact conformation characterized by smaller distances 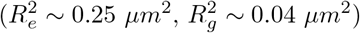.

This transition to a more compact state by increasing *L*, hereafter referred to as L-transition, is quantified in Fig. 1C, where the deviation from the reference value is indicated for different domain sizes *L*_*d*_. We indeed observe a significant deviation compared to the reference case at small *L* ≲ 1Mbp of about

50 *‐* 60% which drops to 5% for *L* ≳ 2Mbp (for *L*_*d*_ = 292kbp). For all *L*_*d*_ values, there is a transition almost at the same *L* (∼ 1 *‐* 2Mbp). The amplitude of the L-transition however depends on *L*_*d*_, with larger domains exhibiting significantly more expanded conformations upon reducing *L*.

To further illustrate the variation in structural features when *L* varies, we plot the mean squared internal distance *R*^2^(*s*) between two loci inside the domain as a function of their genomic distance *s* (*s* = |*j ‐ i*|) (Fig. 1D) in the (pseudo) stationary regime. We observe that the transition between the low and high *L*-regimes is particularly significant at large genomic distance (*s* ≳ 50 kbp). This is consistent with the previous observation that larger domains are more impacted by *L*.

#### 3.1.2 Impact on mobility

In this section, we study how *L* may impact the mobility of monomers inside the domain d and more generally the dynamical properties of the region for *L*_*d*_ = 160kbp. For this, we monitor the average mean squared displacement (MSD) of individual monomers within the domain (*g*_1_(Δ*t*), Fig. 2A), the MSD of the center of mass of the domain (*g*_3_(Δ*t*), Fig. 2B) and the MSD of its end-to-end vector (Fig. 2C). Note that, these three quantities reach a steady-state very rapidly (*≈* 10^7^ MCS) regardless of the value of *L*.

**Figure 2.**
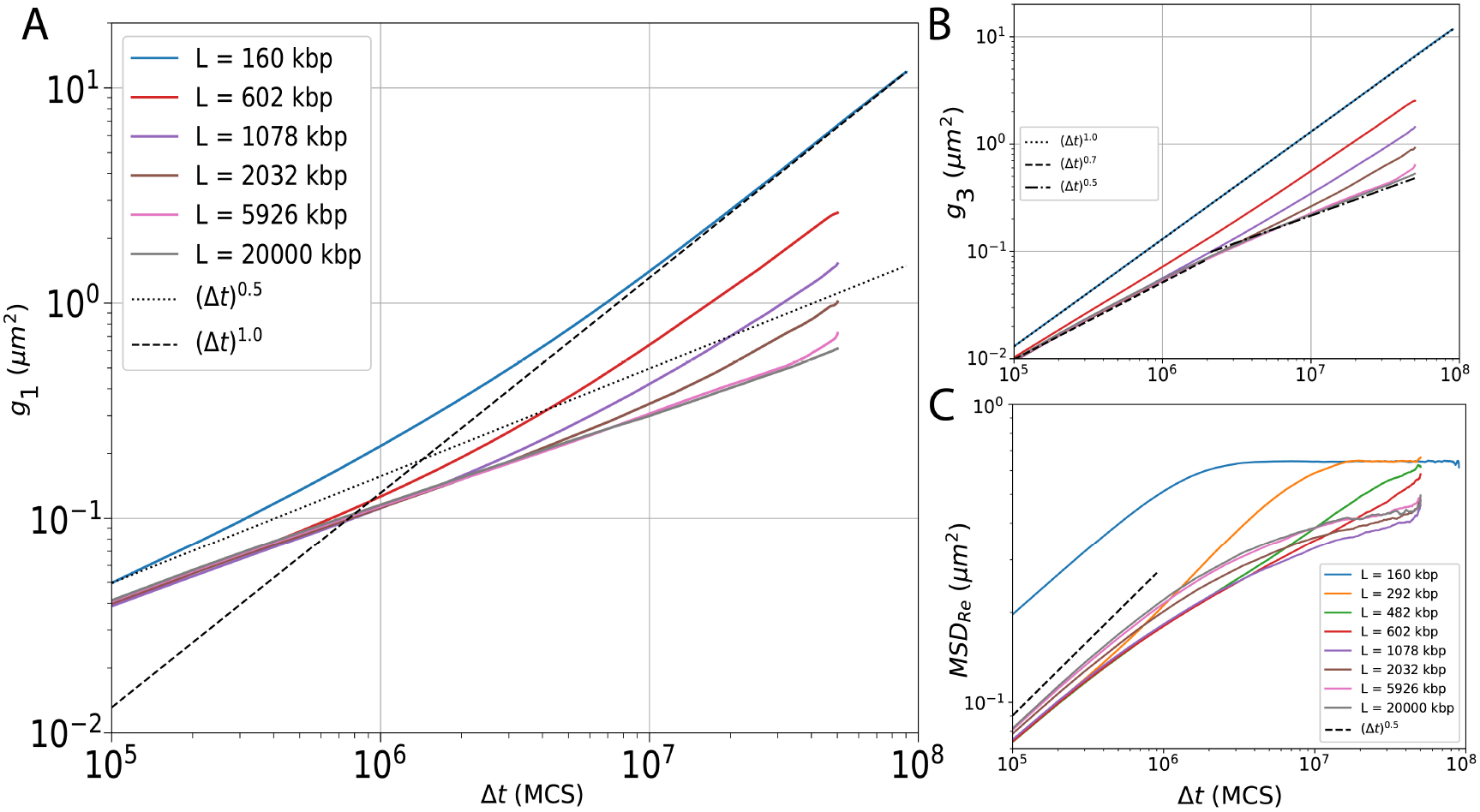
Mean squared displacement (MSD) as a function of time-lag Δ*t* for: (A) Individual monomer MSD inside the domain *g*_1_(Δ*t*). (B) Center of mass MSD of the domain *g*_3_(Δ*t*). (C) MSD of end to end vector of the domain. (A-C) In all simulations volumic fraction ϕ = 0.055 was kept constant.

**Figure 3.**
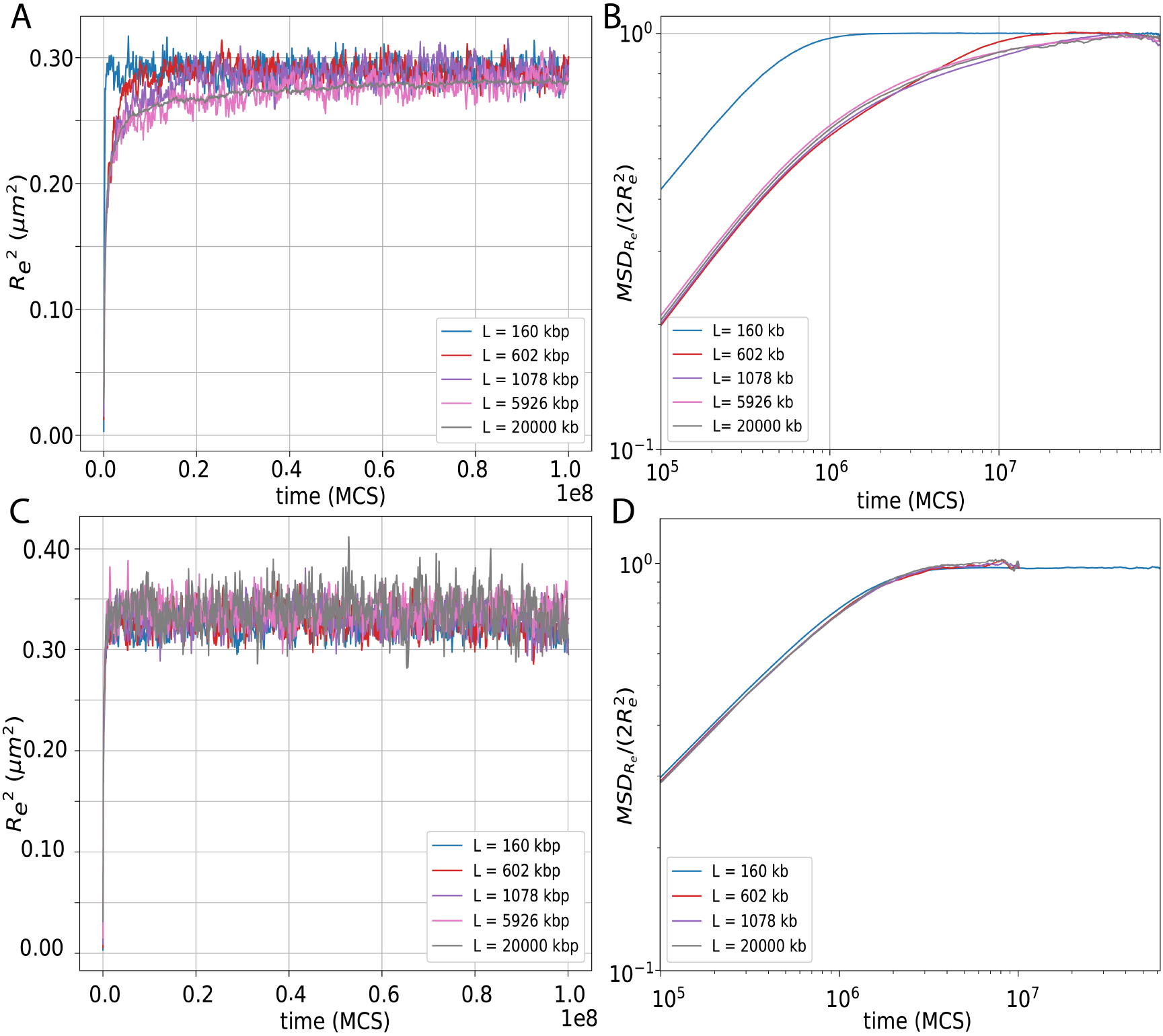
(A) Time evolution of a 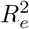 of 160kbp domain for different total polymer length *L*, in absence of excluded volume interactions. (B) Time evolution of MSD of end to end vector of the domain (normalized by the plateau value) for different polymer length *L*, without excluded volume interactions. (C) Time evolution of a 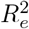 of *L*_*d*_ = 160kbp domain for different total polymer length *L* where domain *L*_*d*_ is disconnected from the polymer chain of length *L ‐ L*_*d*_. (D) Time evolution of MSD of end to end vector of the domain (normalized by the plateau value) for different polymer length *L*, for the case same as (C). (A-D) In all simulations volumic fraction ϕ = 0.055 was kept constant.

When the size of the full chain equals the size of the domain (*L* = *L*_*d*_, blue curves in Fig. 2), we observe the behavior of a simple Rouse polymer [38]. The center of mass undergoes normal Brownian motion with *g*_3_(Δ*t*) *≈* DΔ*t*. Individual monomers exhibit a crossover between a sub-diffusive regime (*g*_1_(Δ*t*) *∝* Δ*t*^1*/*2^) at small timescales and a diffusive behavior at larger timescales, which describes the collective motion of the monomers with the center of mass of the whole polymer.

When *L* increases, for *g*_3_ (Fig. 2B), we observe that the motion of the center of mass depends on *L*, with larger *L* leading to smaller MSD for the same time-lag. Indeed, the global motion of the domain is impacted by the rest of the chain, longer chains having slower diffusion coefficients [38]. Interestingly, beyond the structural L-transition described above (*L* ≳ 2Mbp), *g*_3_ does not significantly depend on *L*, at least on time-scales where the motion of the center of mass of the full chain remains negligible (Δ*t* ≲ 10^9^ MCS). In this case, *g*_3_ of the domain is sub-diffusive (*g*_3_ *∝* Δ*t*^0.7^ for Δ*t* ≲ 2.10^6^ MCS and *g*_3_ *∝* Δ*t*^0.5^ for Δ*t* ≳ 2.10^6^ MCS) as expected for a sub-chain embedded into a larger polymer [32]. For *g*_1_ (Fig. 2A), we observe that at small timescales, the monomer mobility is Rousean (*g*_1_(Δ*t*) *∝* Δ*t*^0.5^) and does not depend on *L*. At larger timescales, *L* starts to impact *g*_1_ as crossover towards the motion of the center of mass of the domain (*g*_3_) is occurring. Again, for *L* ≳ 2Mbp, *g*_1_ becomes relatively independent of *L*.

The MSD of the end to end vector of the domain (Fig. 2C) quantifies the relative mobility of the monomers within the domain and may characterize the motion between regulatory elements like promoters and enhancers [39]. For all *L* values, such MSD is characterized by a sub-diffusive regime (MSD*∝* Δ*t*^0.5^) at small time lags Δ*t* that crosses over to a plateau 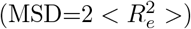 for larger time lags when configurations at time *t* and *t* + Δ*t* are fully decorrelated. However, the equilibration time required to reach this plateau sharply increases with increasing *L*, which can be interpreted as follows. The monomers near the ends of a polymer have more degree of freedom (or are subject to fewer constraints). Therefore, when *L* = *L*_*d*_, maximum mobility is reached (blue curve, Fig. 2C) as expected. For *L* > *L*_*d*_, the MSD is slowed down and it takes much longer time to reach the plateau. Interestingly, we observe a similar behavior as for the equilibration time for *R*^2^ (Fig. 1B): for *L* ≲ 1 *‐* 2Mbp, the decorrelation time (time to reach the plateau) is an increasing function of *L*; for *L* ≳ 1 *‐* 2Mbp, MSD reaches a slightly rising (pseudo) stationary state, similar for every *L*. Intriguingly, for *L* values beyond the structural L-transition where we see higher domain compaction (*L* ≳ 2Mbp), the relative mobility of the ends of the domain is weakly but significantly increased (pink and brown curves in Fig. 2C), indicating that the variation in structural and dynamical properties of the domain may be coupled: slightly more compact organization leading to slightly faster relative diffusion dynamics of the region extremities.

To conclude, the reference dynamical and structural properties for a given *L*_*d*_ (i.e., those obtained for *L* = *L*_ref_) may be fully captured by simulations of smaller polymer chains, providing that *L* is beyond the L-transition (*L* ≳ 2 Mbp in our model).

### 3.2 Excluded volume interaction, chain connectivity and topological constraints are necessary for the L-transition

In order to dissect the causes of the L-transition, we analyze the role of excluded volume and chain properties.

#### 3.2.1 Impact of excluded volume and chain connectivity

In simple polymer models, like Gaussian chains [38], where excluded volume is neglected, the (steady-state) statistical properties of a subchain should not depend on the rest of the chain (i.e. of *L*). In Fig.3A, we indeed verify that, if we remove the excluded volume constraint in our simulations, 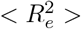 is independent of *L*. Moreover, we also observe that the internal dynamics is also conserved (Fig. 3B).

To further investigate the contribution of chain properties while preserving excluded volume interactions and volumic fraction, we disconnect the domain of length *L*_*d*_ from the rest of the polymer (see Models and Methods). Effectively, the system is now comprised of two polymer chains, one of length *L*_*d*_ and another of length *L ‐ L*_*d*_. As done previously, we systematically vary *L* to study the effect on the structure and dynamics of the domain. In this case, we observe (Fig. 3C,D) that the structural and dynamical properties remain conserved and is independent of the polymer length *L*. This suggests that the L-transition observed previously is invariably related to the embedding of the subchain inside the larger polymer.

#### 3.2.2 Impact of topological constraints

Long confined polymers like chromosomes [40] whose sub-segments generally cannot cross one another (like a macroscopic string) are subject to topological constraints which may considerably affect the polymer conformation and dynamics [41]. The importance of such constraints depends on the ratio between the contour length of the polymer *L*_*c*_ = *Nb* and the so called entanglement length, *L*_*e*_ [42], which can be phenomenologically estimated via the relation [43]:

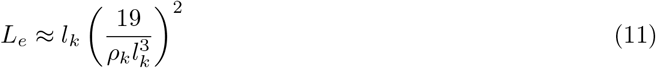

with ρ_*k*_ = (ρ_*bp*_/n)(b/*L*_*k*_) the volumic density in Kuhn segments, and *L*_*k*_ the Kuhn length characterizing the rigidity of the chain and depending on the bending energy *k*_*int*_ [17]. *L*_*e*_ could be related with the tube diameter in the reptation model [38] or to the crossover time between a Rouse-like motion and a reptation-like motion [42]. With the model parameters investigated above (*L*_*k*_ = 100nm and ρ_*bp*_ = 0.009bp/nm^3^), we have *L*_*e*_ = 10.9μm *≡* 556 kbp (see Table S1). If *L ≫ L*_*e*_, topological constraints are strong and the large-scale dynamics of the polymer might be very slow, meaning that the characteristic timescales of equilibration (or of decorrelation) of the lower Rouse modes of the chain (i.e., corresponding to large-scale variables, like the end-to-end vector) could be very large, and that the polymer keeps memory of the initial topological state of the chain (eg, presence or not of knots, large-scale organization) for a very long time [17, 44], larger that the total simulation time (that corresponds approximately to ∼ 1 hour of real time).

Thus, topological constraints that implicitly depend on excluded volume and confinement might be the main driver of the L-transition.

##### Strand crossing affects the L-transition

First, we directly investigate the role of topological constraints on the domain structure and dynamics by introducing topology changing, strand crossing moves following the procedure described in the sec. 2.4. Such moves, biologically motivated by the putative activity of topoisomerase II *in vivo*, [45, 46], allow chain crossing with a low frequency without otherwise affecting the excluded volume properties.

We observe that the convergence towards a stationary state for the structural properties of a domain is much faster and becomes independent of the total polymer length (Fig. 4A). It is to be noted that there is no L-transition anymore which conclusively confirms our hypothesis that the L-transition observed in the absence of strand crossing moves is driven by topological constraints. This is also reflected in the dynamics, relative mobility of the monomers within the domain is the same irrespective of the total polymer length and the decorrelation time (time to reach the plateau) remains the same, independent of *L* (Fig. 4B).

**Figure 4.**
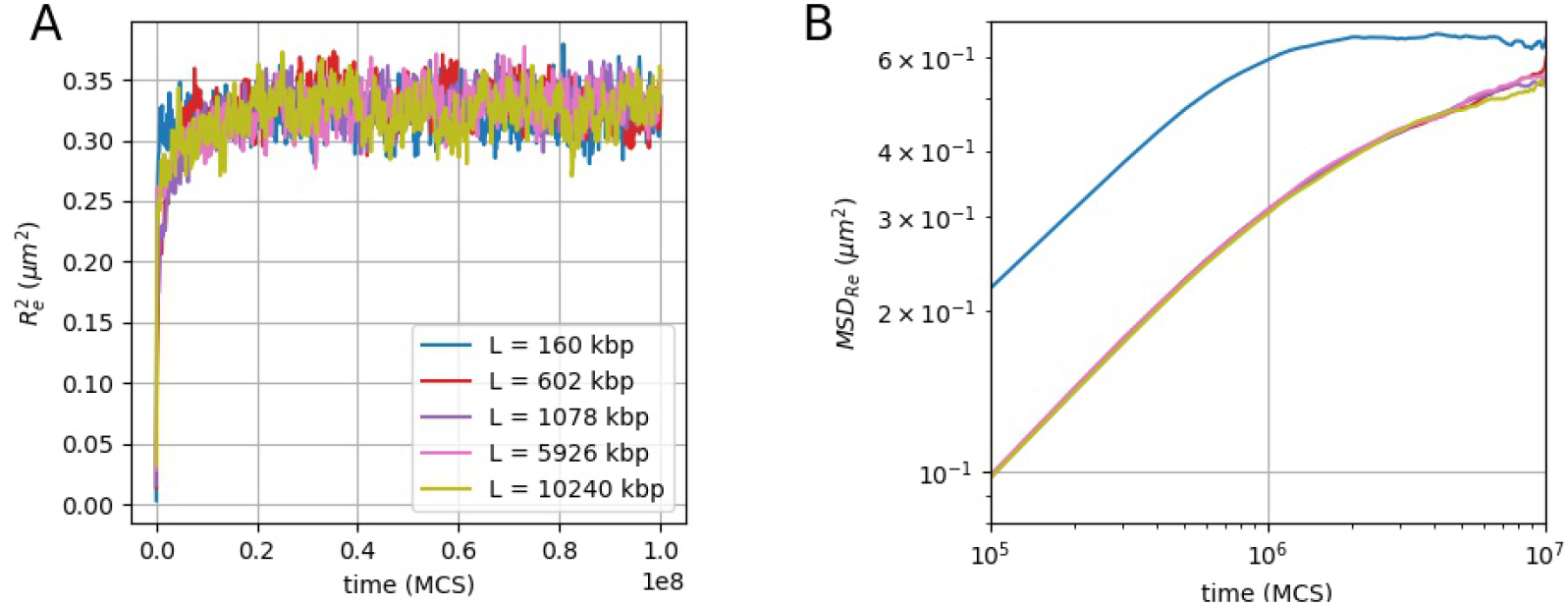
Topology changing moves: (A) Deviation of 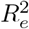 of 160kbp domain from the reference as a function of *L*. (B) Time evolution of MSD of end to end vector of the domain for different polymer length *L*. (A,B) In all simulations volumic fraction ϕ = 0.055 was kept constant.

We can also confirm that the weak increase in mobility observed in the absence of topology changing moves (Fig. 2C) is directly linked to the transition seen in structural properties and is driven by topological constraints.

##### Impact of entanglement length

In our simulations, as we increase *L*, the system transitions from a regime of weak topological constraints (*L*/*L*_*e*_ ≲ 1) to a regime of strong constraints (*L*/*L*_*e*_ *≫* 1). We analyze the properties of the end-to-end vector of the domain d (*L*_*d*_ = 160kbp) for polymer models with different entanglement lengths *L*_*e*_. In addition to the default one 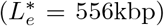 studied above, we explore 3 other situations with lower *L*_*e*_ by varying the dimensions of the simulation box, i.e. modifying the monomer volumic density and thus *L*_*e*_ [17]: 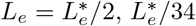 and 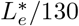 (see Table S1). For each situation, we also reach a (pseudo)-steady-state (Fig. 5A) that allows us to precisely define 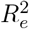. In Fig. 5B, we observe that the L-transition is also present for the other *L*_*e*_ values. Importantly, the amplitude of the transition is larger for lower *L*_*e*_ and the critical value of *L* where the system shifts from less to more compact is smaller for lower *L*_*e*_ values. We also observe that, not only the end-to-end distance, but all internal distances are affected by *L*_*e*_ (Fig. S5). The differences seen at large genomic distances between *L* = *L*_*d*_ and *L* = *L*_ref_ are stronger for smaller *L*_*e*_. In this case, the two curves also start to deviate from each other at shorter genomic distances, meaning that topological effects become more prominent at smaller genomic scale for smaller *L*_*e*_. All this confirms that the L-transition is driven by a change of topological regime for the whole chain, and that large domains (or genomic distance) compared to the entanglement length (*L*_*d*_/*L*_*e*_, *s*/*L*_*e*_ ≳ 1) would be impacted more (red curve, Fig. 1C).

**Figure 5.**
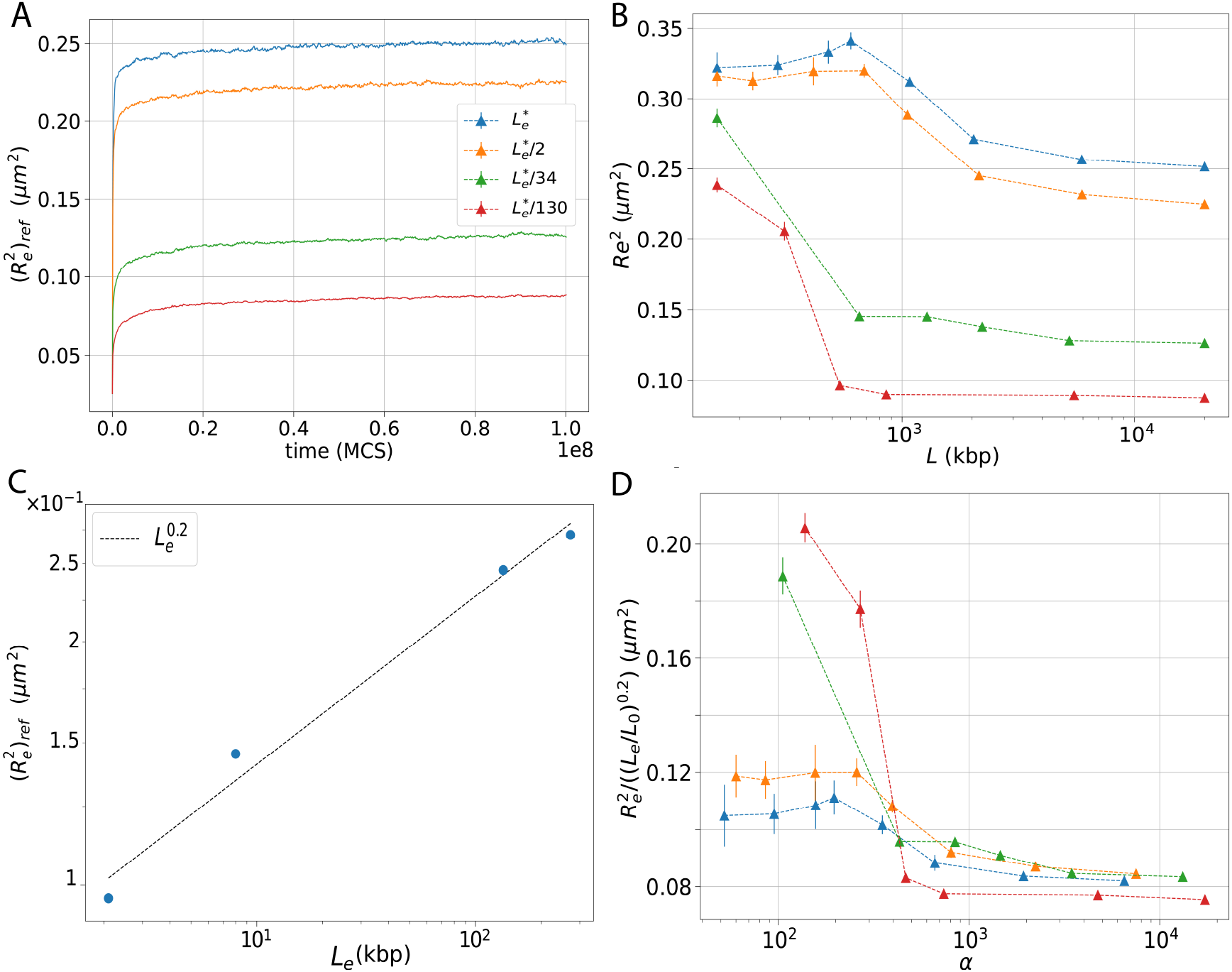
(A) The reference end-to-end distance squared as a function of time for different entanglement lengths *L*_*e*_. (B) 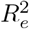 of 160 kbp domain as a function of total polymer length *L* for different *L*_*e*_ values. For a given *L*_*e*_ value, all the simulation were done at constant volumic fraction (Table S1). (C) The pseudo-steady-state value 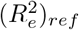 as a function of *L*_*e*_. (D) 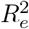 normalized by (*L*_*e*_/*L*_0_)^0.2^ as a function of α, where α = *L*/(*L*_0_(*L*_*e*_/*L*_0_)^0.2^), for polymers of different entanglement length *L*_*e*_. α ≳ 1000 delimits the region where structural properties are independent of *L*.

To quantify this precisely, in Fig. 5C, the pseudo-steady-state value 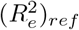 of 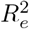 in the reference case (*L* = 20Mbp) is plotted as a function of entanglement length *L*_*e*_, and empirically we obtain a scaling law for the end-to-end distance squared after the transition 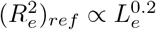. As expected, strong topological constraints lead to more compact domains (i.e., smaller 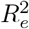). We define α = *L*/(*L*_0_(*L*_*e*_/*L*_0_)^0.2^), a dimensionless quantity to estimate the extent of topological constraints in the system with *L*_0_ = 1kbp, an arbitrary default value. Phenomenologically, we find that α ≳ 1, 000 corresponds to the values of *L* for which the structure of the domain is not affected anymore by increasing *L* (Fig. 5D). In other words, more practically, a polymer of length *L* > *L*_*α*_ *≡* 10^3^ *∗* (*L*_0_(*L*_*e*_/*L*_0_)^0.2^), would preserve the structural properties of the domains, as if it were simulated within the whole (reference) chromosome.

Altogether, our observations suggest that the L-transition is driven by topological constraints. In the regime where topological constraints are relevant (α ≳ 10^3^ or *L* ≳ *L*_*α*_), the subchain maintains its topological state during the whole simulation, i.e., a knot-free topology in our case. This limits the structural fluctuations of the sub-chain and leads to more compact structures, as typically observed for crumpled or ‘fractal’ polymers [41, 44, 47] that are often associated with compact, space-filling curves [48]. Importantly, we have to emphasize here that when the domain size *L*_*d*_ is greater than the entanglement length (*L*_*d*_/*L*_*e*_ *≥* 1), the domain structure is very sensitive to chromosome length when *L ≤ L*_0_ *∗* (*L*_*e*_/*L*_0_)^0.2^. On the contrary when *L*_*d*_ is smaller than the entanglement length, the length of the chromosome being simulated has minimal effect i.e global topological constraints do not drastically affect the structure and organization of the domain (*L*_*d*_/*L*_*e*_ << 1, orange curve in Fig. 1C).

### 3.3 Total polymer length impacts coil-to-globule transition of the subchain

The modeling of genome 3D organization and in particular of the large-scale folding into active and inactive compartments has led to the introduction of copolymer models of chromatin. Beyond the excluded volume constraint, these models [5, 6, 13, 17, 49–51] include self-attractions (*J*) between loci sharing similar epigenomic contents (see Models and Methods). *J* effectively accounts for different physico-chemical contributions such as charge modulated nucleosome-nucleosome interaction [52], architectural proteins bridging [53–55] or solvent quality [56].As a universal behaviour, a homopolymer with self-attraction *J* acting on all its monomers undergoes a coil-globule transition from an expanded coil-like conformation for |*J*| < |*J*_*c*_| to a compact globular conformation for |*J*| > |*J*_*c*_| [57, 58], where *J*_*c*_ is the transition point (called the “theta” point). Such collapse transition depends on the (finite) size of the chain [15, 16] with sharper transitions for larger lengths of the self-interacting chain.

In this section, we employ a standard copolymer model to study how the total polymer length *L* would affect the coil-globule transition of a particular region of interest, taking as reference the situation for *L*_ref_ = 20*Mbp*. To simplify, we consider a region of size *L*_*d*_ = 160 kb of a particular epigenomic state embedded into a neutral chain of size *L*. For all *L*, as we vary the strength of self-attraction *J*, the subchain gradually collapses (Fig. 6) with 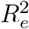 of the region following a standard coil-globule transition, starting from an initial expanded state (higher 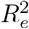) to a more compact (lower 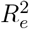). Beyond the theta-collapse, in the globular state, 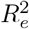 is independent of *L*. However, for |*J*| < 0.2*k*_*b*_*T*, during the collapse, 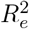 becomes more sensitive to *L*. Consistent with our observations in the pure self-avoiding case that larger *L* lead to more compact subdomains (Fig. 1A), larger *L* would be more strongly collapsed for a given *J* value. Interestingly, the coil-globule transition curves are almost similar for all *L* ≳ *L*_*α*_ *≈* 2Mbp that corresponds to the critical *L* value of the L-transition observed in the pure self-avoiding system.

**Figure 6.**
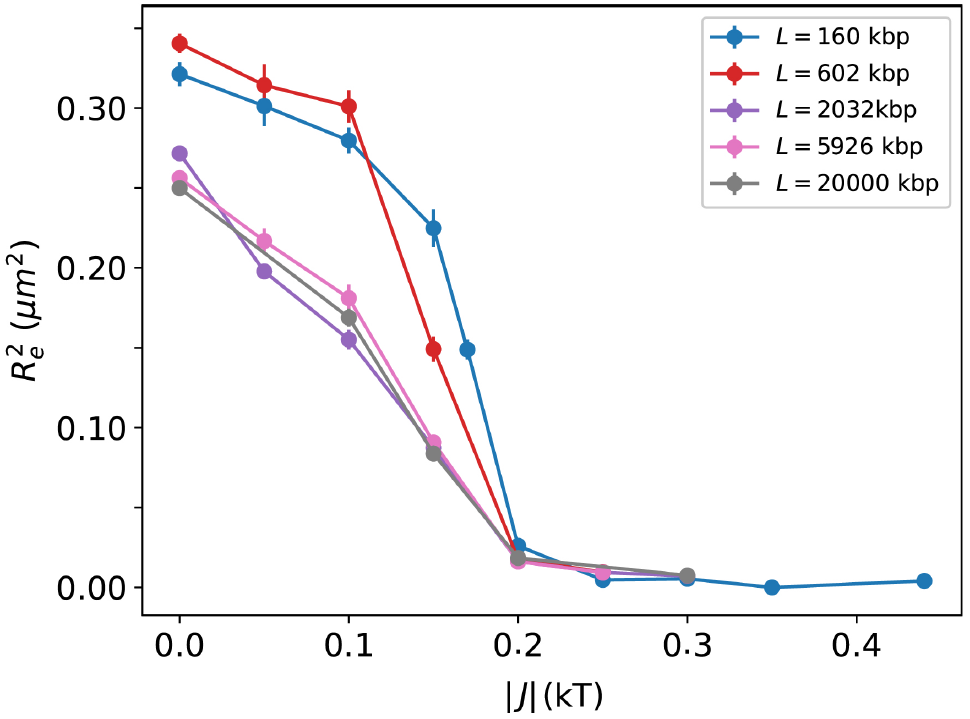
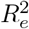 for a 160 kb region as a function of interaction strength |*J*| for different total polymer lengths *L*. In all simulations volumic fraction ϕ = 0.055 was kept constant.

**Figure 7.**
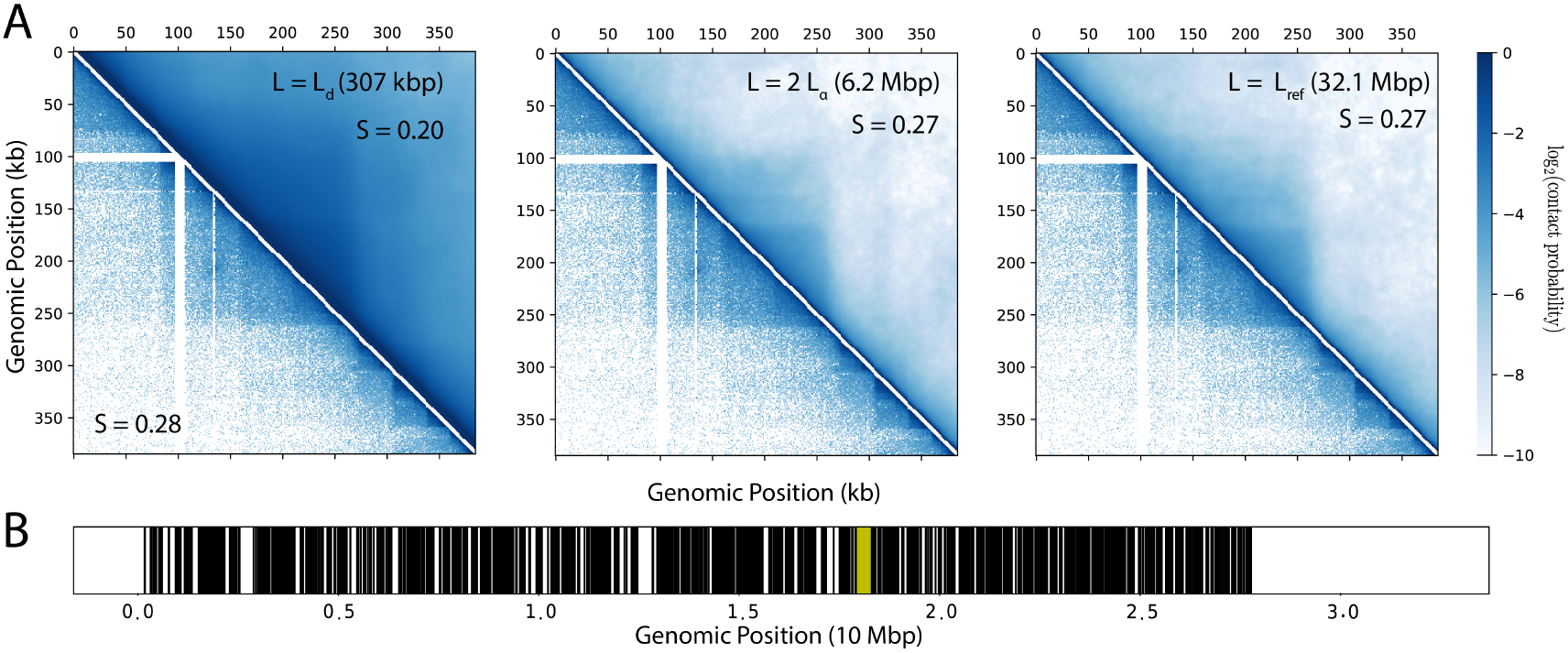
Simulated (upper part) and experimental [30] (lower part) Hi-Cs of a Drosophila domain for simulations with (Left) *L* = *L*_*d*_ = 307kbp and *J* = *‐*0.17*k*_*b*_*T*. (Middle) *L* = 6.2 Mbp and *J* = *‐*0.135*k*_*b*_*T*. (Right) *L* = 32.1Mbp and *J* = *‐*0.135*k*_*b*_*T*. It is to be noted that the 3D organization of the domain is very similar in the case of *L* = 6.2Mbp and *L* = 32.1Mbp. The saddle-strength compartment score values *S* are shown for each case. The monomers of the interacting epigenomic state (M) and the non-interacting U-state are shown at the bottom. In all simulations volumic fraction ϕ = 0.028 was kept constant. (B) Shows the distribution of black domains along chromosome 3R, the domain we have considered is highlighted in yellow. *Data from [60]*.

While, when the sub-chain is collapsed, the effect of *L* is not significant, it plays a crucial role in the structure and dynamics of the domain in partially collapsed chain, in particular in the case of systems where strong topological constraints *L*_ref_ /*L*_*e*_ >> 1 are present in the reference case, as in the genomes of Drosophila (*L*_ref_ /*L*_*e*_ *≈* 70 [17]) and higher eukaryotes. In this case, our analysis shows that topological constraints may be needed to be accounted for when α(*L*_ref_) > 10^3^, in particular when the region is not fully collapsed or globular (see Fig.6) as often observed in biological context [17]. In organisms with long chromosomes (α(*L*_ref_) *≫* 10^3^) as considered above, we expect that the minimal length of polymer to simulate to preserve the structural reference properties (as in the full chromosome), is *L* ≳ *L*_*α*_ *≈* 2 *‐* 4 Mbp. In the case of short, reference chromosomes (*L*_ref_ < *L*_*α*_) like yeast, comprised of smaller chromosomes (*L*_*e*_ = 2.2 Mbp, *L*_*α*_ *≈* 5 Mbp), α(*L*_ref_) = 236 *≪* 10^3^, we expect topological constraints not to be too relevant and thus that *L* has only a limited impact (Fig. S6, S7).

### 3.4 Total polymer length and epigenomic context drive genome folding

In the previous sections, we have not accounted for the extrinsic contributions which could arise (e.g.) from the putative epigenomic interactions between the domain of interest and the rest of the genome, and which could potentially also affect the local structure of the region. We have considered regions which are completely isolated illustrating how topological constraints impact the 3D organization of the region. But it is evident that the overall epigenomic context and associated interactions would be pertinent to fully describe the structure and dynamics of the domain [59].

To understand how coupling the effect of topological constraints (total polymer length *L*) with the environmental epigenomic context might affect the 3D organization of the domain of interest, we simulate a domain corresponding to the region 17, 955 to 18, 262 kbp in chromosome 3R of Drosophila (*L*_*d*_ = 307 kbp), which is denoted as a ‘black’ epigenomic domain (silent chromatin) by the classification done in [60]. In its native context, this domain is embedded within a polymer of size *L*_ref_ = 32.1Mbp and the domain can interact with all the other black domains along the chromosome.

We simulate three scenarios: (i) *L* = *L*_*d*_; (ii) *L* = 6.2 Mbp, a length (*≈* 2 *∗ L*_*α*_) that should capture the polymeric, reference case based on our theoretical analysis; and (iii) *L* = *L*_ref_. For each case, we vary *J* and compute Hi-C maps for different radius of capture *R*_*c*_, and estimate the (*J, R*_*c*_) parameters that best reproduces the experimental HiC of the region in the each scenario by minimizing a χ^2^-like score (see Models and Methods, Fig.S8). In contrast to our previous analysis, we now are considering a domain that is in a dense interacting environment, potentially also subject to attractive interactions with domains of similar epigenomic state along the chain.

For the three cases, the best fitting radius of capture is *R*_*c*_ = 60nm. Interestingly, we find two different optimal *J* values (*J* = *‐*0.17*k*_*b*_*T* for scenario *i* and = *‐*0.135*k*_*b*_*T* for scenario *ii* and *iii*), confirming that simulating *L* = *L*_*d*_ and *L* = 32.1Mbp is quantitatively different also for a highly interacting domain embedded inside a long chromosome with distributed interacting regions. The degree of agreement with the experimental Hi-C map is further found to be slightly better for simulations including longer flanking segments (Fig. 7A) with a Pearson correlation between predictions and experiments of 0.87 for *L* = 2*L*_*α*_ and *L*_ref_ versus 0.86 for *L* = *L*_*d*_. Consistently, the saddle-strength compartment score *S*, a measure of compartmentalization (see Model and Methods), is found to be significantly better reproduced by simulations with longer flanking regions, with the predicted value *S* = 0.27 for *L* = 2*L*_*α*_ and *L*_ref_ closely matching the experimental value *S* = 0.28 — to be compared with the lower value *S* = 0.20 obtained for *L* = *L*_*d*_, using the best-fit (*J, R*_*c*_) parameter combination in each case.

We note that changing the total polymer length significantly alters the 3D organization of the region of interest. Interestingly, we find that the length scale predicted by our analysis on topological constraints is sufficient to reproduce the folded structure of the domain as reported in the experimental map (Fig. 7A, middle). It is important to consider large-enough polymer lengths to correctly predict the conformational statistics of individual embedded domains, which requires a proper account of the effects of the topological constraints imposed by the surrounding flanking regions.

## 4 Discussion and conclusion

In this paper, we have illustrated the role of total polymer length on the structural and dynamical properties of a subchain in the context of 3D genomics. Our analysis gives quantitative predictions on the minimal length of chromatin to be simulated around a specific region of interest to capture correctly polymeric properties of the region, such as spatial distances, contact probabilities or mean-squared displacements that are accessible experimentally.

Using a simple self-avoiding homopolymer, we have first systematically studied how increasing the total polymer length *L* at fixed volumic fraction would affect the 3D organization of an embedded region of arbitrary size (Fig. 1), taking the case of a long chromosome (*L* = 20 Mb) as a reference. For short polymers (*L ≤* 2 Mb), the internal structure of the subchain is more expanded and equilibrates much faster. For longer polymers (*L ≥* 2 Mb), simulations indicates the presence of a very slowly expanding, more compact, (pseudo)steady-state, characteristic of crumpled polymers [29, 41]. The dynamics of the subchain exhibits also differences in the local mobility of monomers (Fig. 2) when the total polymer length is varied, although to a lesser extent than the effect on structure.

We have observed that without excluded volume interactions and chain connectivity and with strand crossing, the structure and dynamics of the domain is not affected anymore by the total polymer length (Fig. 3, 4), demonstrating that topological constraints are responsible for the different behaviors observed when varying *L*. Furthermore, we quantified the extent of topological constraints to be considered by a single parameter (α) that depends on the entanglement length *L*_*e*_ and thus on chromatin fiber stiffness and volumic density. This parameter helps in estimating the minimal length *L*_*α*_ of chromatin to be simulated to capture the correct conformational and dynamical properties (Fig. 5). These results would significantly help in extracting accurate physical measures of different domains using simple polymer models [13], as well as in providing a convenient empirical criterion for the minimal system size one must employ in order to accurately describe single-domain folding structures with minimal computational effort.

We have also used a simple copolymer framework to understand how the total polymer length may affect the folding properties of the domain of interest. Beyond the Θ collapse, we showed that the globule-state is unaffected by the total polymer length (Fig. 6). However, before the collapse (a biologically-relevant regime [5, 6, 17]), we found that *L* may impact the folding and should be chosen carefully like in the pure self-avoiding case.

In the last part of our results, we explored how accounting for the epigenomic environment may impact our conclusions by contextualizing the model to experimental data: Drosophila [30] (with strong topological constraints) and yeast [31] (weakly constrained). The correct behaviour of a domain in a dense interacting environment seems to be correctly captured if *L* is large enough to account for the polymeric behavior (Fig. 7). In the Drosophila case, *L* may alter the 3D organization of the region (Fig. 7) while in the yeast case, it doesn’t play a crucial role (Fig. S6). This conclusion is further compounded by the fact that the interacting domains (which mostly corresponds to genes [7]) in yeast are very small (< 10kbp) and thus topogical effects have less impact on the organization (Fig. 1C). It is to be noted that this might depend on the specific system under consideration. Beyond the epigenomic context, it is also likely that association with nuclear lamina [61] or other nuclear bodies [62] or other architectural mechanisms such a SMC-mediated loop extrusion [4] may potentially provide additional geometric and physical constraints that may contribute to the behavior of a domain of interest and that may depend on the size *L* of the simulated polymer.

Overall, our results suggest that the intrinsic and extrinsic environmental contribution acting on a subchain of interest are well captured by simulating a total polymer length *L* such that *L* ≳ *L*_*α*_ in the case where the reference chromosome is long and the domain of interest is coiled or partially-collapsed. Conversely, one may neglect the contribution of the flanking regions, and hence restrict the simulations to the sole domain of interest *L ≈ L*_*d*_ if either of the following conditions are met: (i) the domain is strongly collapsed, (ii) the domain is sufficiently short (*L*_*d*_ *≪ L*_*e*_), or (iii) the corresponding reference chromosome is sufficiently short (*L*_ref_ *≪ L*_*α*_). Such conclusions should be broadly applicable to quantitative simulations of individual genes and topologically-associated domains (TADs) based on finite-sized polymer systems, which have been employed in a variety of biological contexts. It is to be noted that we focus our analysis mostly on observables capturing the structural and dynamical properties of individual monomers or of pair of monomers within *L*_*d*_. It is thus possible that collective or higher-order quantities may not be well captured even if *L* ≳ *L*_*α*_.

One key result of this work is the requirement of maintaining the system in the correct topological regime to ensure that polymeric properties are conserved. However, this result is based on the physical hypothesis of the model that the polymer strands cannot cross each other (or at least only very rarely), thus maintaining the system in the same topological state during the simulations of long polymers [29]. Experimentally, it was estimated that chromosomes have no or only few knot-like entanglements [26, 27] during G1. However, it is still unclear whether, *in vivo*, such an “unknotted” state emerges solely from the decompaction of the chromosomes from a “knot-free”, mitotic-like initial configuration that would remain devoid of knot-like entanglements thanks to rare strand crossing events [28, 29]. Indeed, the polymer topological state may also be affected by the Topoisomerase II activity [64], putatively in combination of active loop extruding factors [63] like cohesins or condensins which are thought to play crucial role in TAD formation [4]. In our framework, we have introduced topology changing, strand crossing moves in the model mimicking a Topo II-like activity. Our results suggest that such moves may serve to limit the impact of the L-transition reported here (Fig. 4) by reducing the topological constraints. It would thus be very interesting to introduce strand crossing and modulate its propensity in a loop extrusion polymer model [65] to investigate how our conclusions may be modified by these additional contributions.

## Supplementary Information

Additional simulation information and results including Supplementary Table S1 and Supplementary Figures S1 to S9.

## Acknowledgements

We are grateful to Maria Barbi, Jean-Marc Victor, Angelo Rosa and the members of Jost lab for fruitful discussions. We acknowledge Agence Nationale de la Recherche [ANR-18-CE45-0022-01 to A.Z.A., D.J., ANR-21-CE45-0011-01 to D.J., C.V.] for funding. We thank PSMN (Pôle Scientifique de Modélisation Numérique) of the ENS de Lyon for computing resources.

