## Supplementary material for "Topological constraints and finite-size effects in quantitative polymer models of chromatin organization": Table S1 and Fig.S1 to S9

**Content:**

- Table S1
- Figures S1 to S9.

**Table S 1.** Different polymer and lattice parameters used in the different sections of the main text.

| **In Figs.1-6 & S1-S5**: $n$ = 1kbp, $b$=20nm, $k_{int} = 3.217 k_b T$ | | | | |
|---|---|---|---|---|
| $L$ (kbp) | $N$ | $S$ | $\phi$ | $L_e$ (kbp) |
| 75 | 75 | 7 | 0.0547 | 556 |
| 160 | 160 | 9 | 0.0549 | 556 |
| 292 | 292 | 11 | 0.0548 | 556 |
| 482 | 482 | 13 | 0.0548 | 556 |
| 602 | 602 | 14 | 0.0548 | 556 |
| 1,078 | 1,078 | 17 | 0.0548 | 556 |
| 2,032 | 2,032 | 21 | 0.0548 | 556 |
| 5,926 | 5,926 | 30 | 0.0549 | 556 |
| 10,240 | 10,240 | 36 | 0.0549 | 556 |
| 20,000 | 20,000 | 45 | 0.0549 | 556 |
| 160 | 160 | 8 | 0/0781 | 278 |
| 416 | 416 | 11 | 0.0781 | 278 |
| 687 | 687 | 13 | 0.0781 | 278 |
| 1055 | 1055 | 15 | 0.0781 | 278 |
| 2143 | 2143 | 19 | 0.0781 | 278 |
| 5492 | 5492 | 26 | 0.0781 | 278 |
| 20,000 | 20,000 | 40 | 0.0781 | 278 |
| 160 | 160 | 5 | 0.32 | 16.35 |
| 655 | 655 | 8 | 0.32 | 16.35 |
| 1280 | 1280 | 10 | 0.32 | 16.35 |
| 2211 | 2211 | 12 | 0.32 | 16.35 |
| 5240 | 5240 | 16 | 0.32 | 16.35 |
| 20,000 | 20,000 | 25 | 0.32 | 16.35 |
| 160 | 160 | 4 | 0.625 | 4.29 |
| 312 | 312 | 5 | 0.624 | 4.29 |
| 540 | 540 | 6 | 0.625 | 4.29 |
| 858 | 858 | 7 | 0.625 | 4.29 |
| 5494 | 5494 | 13 | 0.625 | 4.29 |
| 20,000 | 20,000 | 20 | 0.625 | 4.29 |
| **In Figs.7 & S6-S8**: $n$ = 800bp, $b$=15nm, $k_{int} = 4.05 k_b T$ | | | | |
| $L$ (kbp) | $N$ | $S$ | $\phi$ | $L_e$ (kbp) |
| 45.6 (yeast) | 57 | 11 | 0.0107 | 2190 |
| 1100 (yeast) | 1375 | 32 | 0.0105 | 2190 |
| 307 (Drosophila) | 385 | 15 | 0.0285 | 676 |
| 6200 (Drosophila) | 7750 | 41 | 0.0281 | 676 |
| 32,100 (Drosophila) | 40,125 | 71 | 0.0280 | 676 |

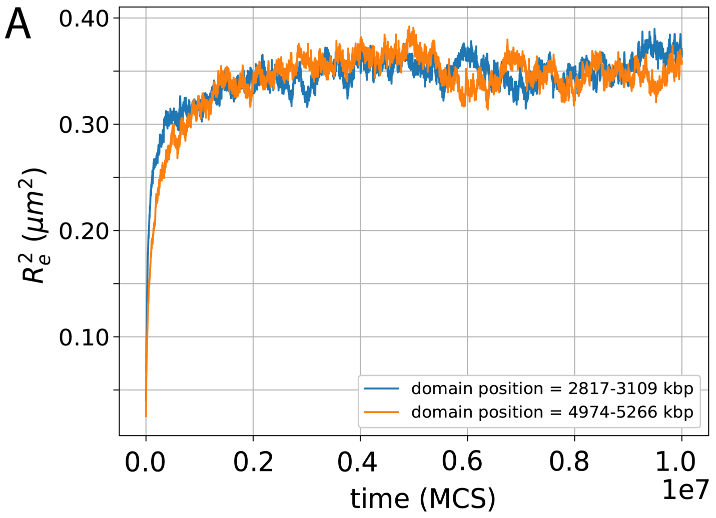


**Fig. S 1.** (A) Evolution of $R_e^2$ of the region of interest (domain) of length $L_d = 292$kbp in a polymer of length $L = 5,926$kbp. The relative position of the domain with respect to the polymer is varied, blue curve, when the domain is in the middle of the polymer and orange, when the domain is shifted towards the right but it is to be noted that the structural variable evolves is the same manner.

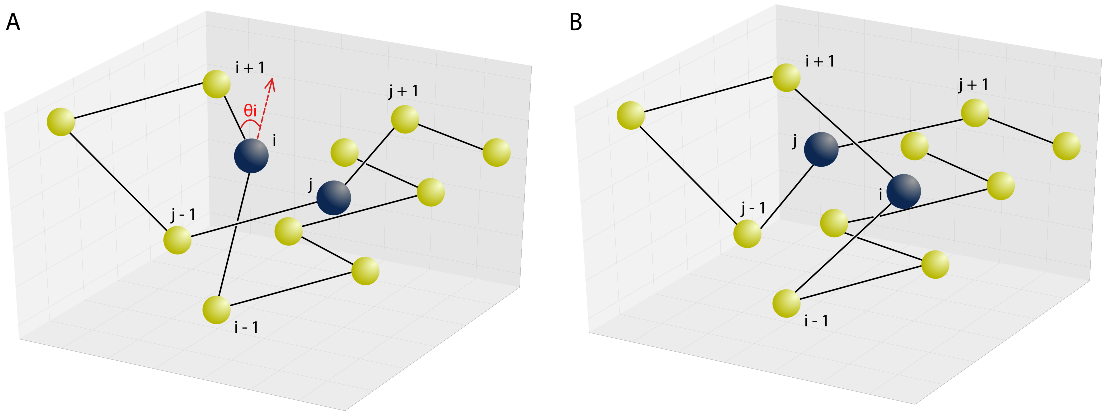


**Fig. S 2.** Illustration of strand crossing moves. (A) Configuration where monomers $i$ and $j$ (in blue) can be swapped such that chain connectivity would be preserved. (B) Configuration after the strand crossing move (swap between position of $i$ and $j$) has been performed.

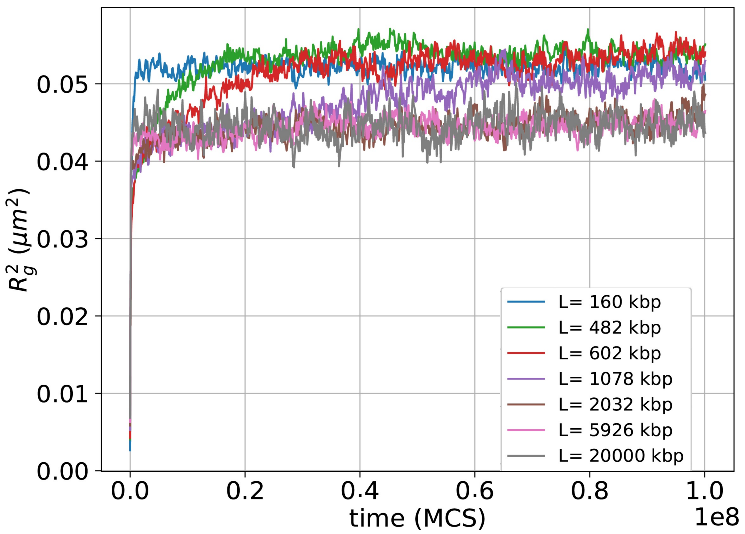


**Fig. S 3.** Time evolution of the squared radius of gyration $R_g^2$ of a $L_d = 160$kbp domain for different total polymer length $L$.

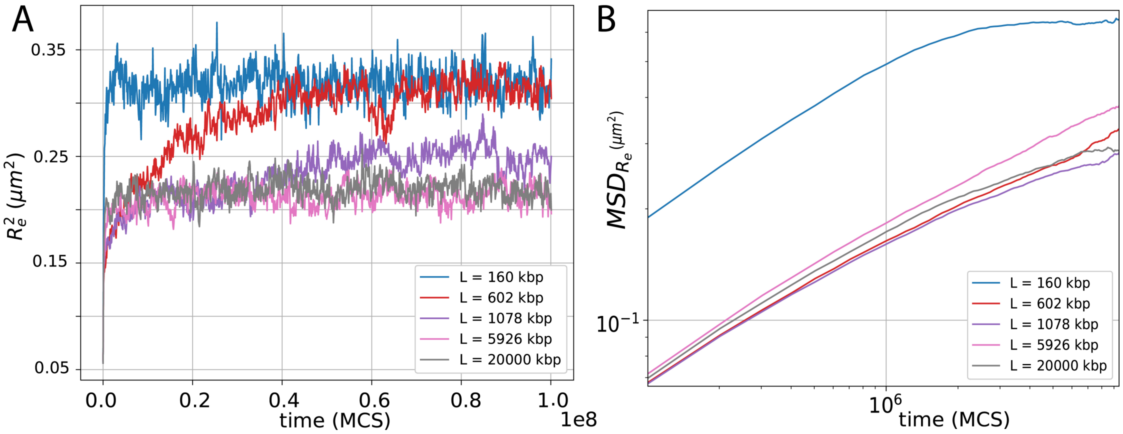


**Fig. S 4.** An L-transition is also observed for initial configurations build by random walk and thus possibly containing knot-like entanglements ("knotted" state). (A) Time evolution of the squared end-to-end distance $R_e^2$ of a $L_d = 160$kbp domain for different total polymer lengths $L$. (B) Mean squared displacement (MSD) as a function of time-lag $\Delta t$ of end to end vector of the domain.

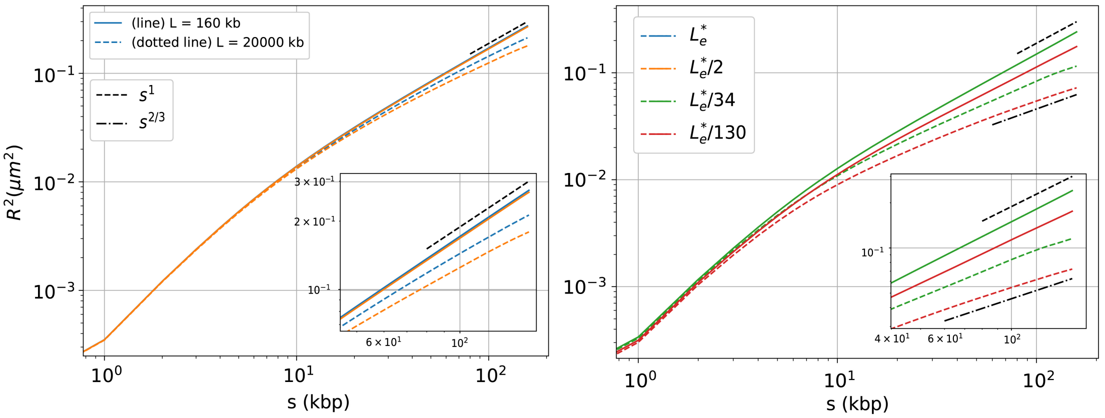


**Fig. S 5.** Average squared 3D distance $R^2(s)$ between monomers of the domain ($L_d = 160$kbp) as a function of genomic distance $s$. Solid lines correspond to $L = L_d$ and dotted lines, $L = L_{ref} = 20,000$kbp. Inset shows a zoom at large genomic distances ($s \gtrsim 50$ kbp) in log-log scale. The four colors indicate four different entanglement lengths with $L_e^* = 556$kbp.

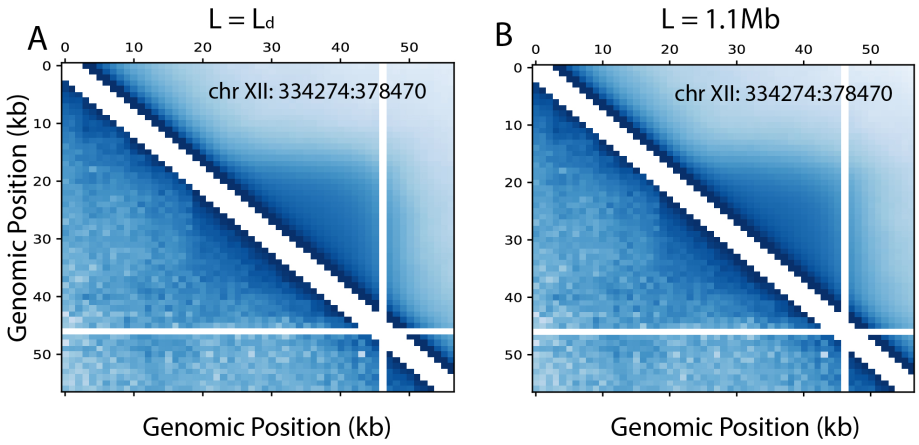


**Fig. S 6.** Simulated (upper part) and experimental (lower part) Hi-C of a yeast domain (chr XII; $334,274 : 378,470$) for simulation with (A) $L = L_d = 45.6$kbp and $J = -0.25k_bT$ (corresponding to Fig.7A); with (B) $L = 1.1$ Mbp and $J = -0.25k_bT$ (Fig. 7B). Pearson correlation value:= 0.92 for $L = 1.1$Mbp and $L = L_d = 45.6$kbp

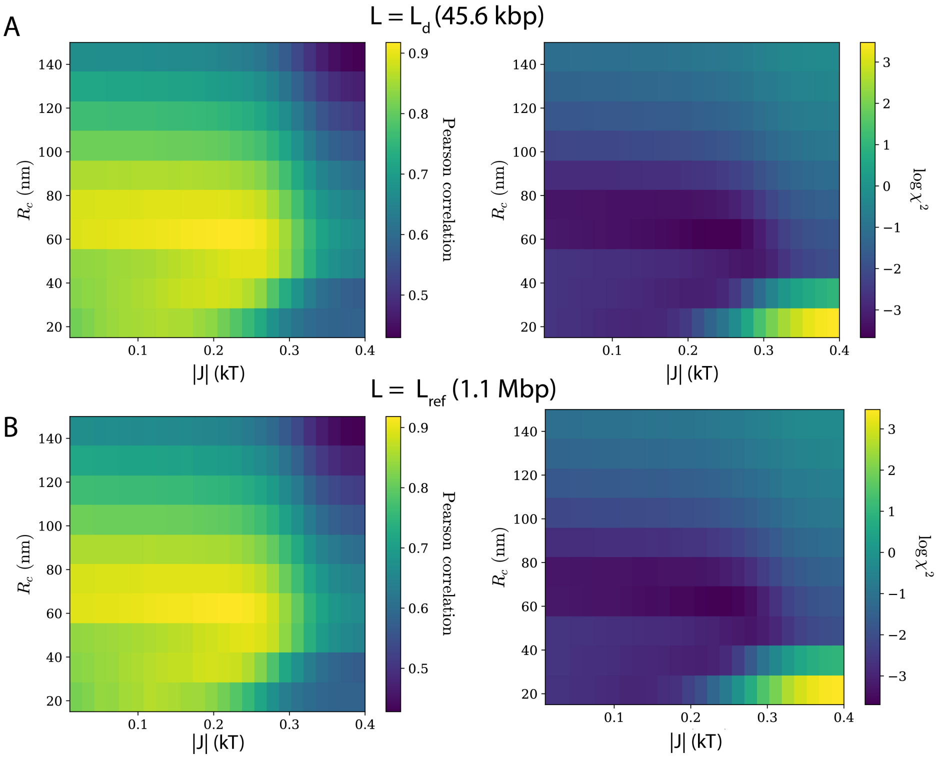


**Fig. S 7. Yeast:** Phase diagram ($R_c$ and $J$) quantifying the predictive power of the simulation relative to experimental Hi-C data using two different metrics: (i) Pearson correlation (left) and (ii) $\chi^2$ as a function of two parameters (right). (A) For $L = L_d = 45.6$kbp, the best fitting parameters (minimum $\chi^2$) being $R_c = 60$nm, $J = -0.25k_bT$ ;(B) For $L = 1.1$Mbp, the best fitting parameters (minimum $\chi^2$) being $R_c = 60$nm, $J = -0.25k_bT$.

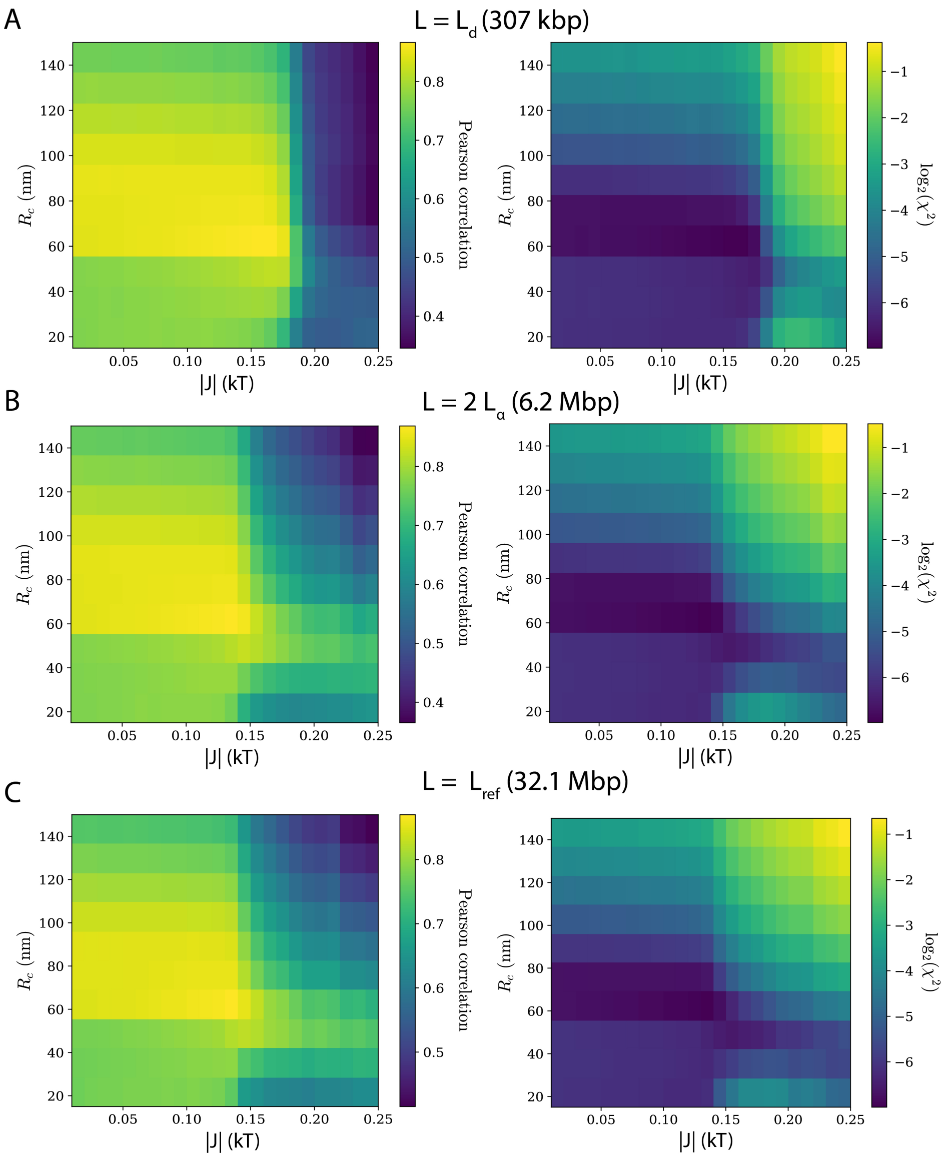


**Fig. S 8. Drosophila:** Phase diagram ($R_c$ and $J$) quantifying the predictive power of the simulation relative to experimental Hi-C data using two different metrics: (i) Pearson correlation (left) and (ii) $\chi^2$ as a function of two parameters (right). (A) For $L = L_d = 307$kbp, the best fitting parameters (minimum of $\chi^2$) being $R_c = 60$nm, $J = -0.17k_bT$ ; (B) For $L \approx 2 * L_\alpha = 6.2$ Mbp the best fitting parameters (minimum of $\chi^2$) being $R_c = 60$nm, $J = -0.135k_bT$ ; (C) for $L = L_{ref} = 32.1$ Mbp, the best fitting parameters (minimum $\chi^2$) being $R_c = 60$nm, $J = -0.135k_bT$. These values are in agreement with maximum Pearson correlation.

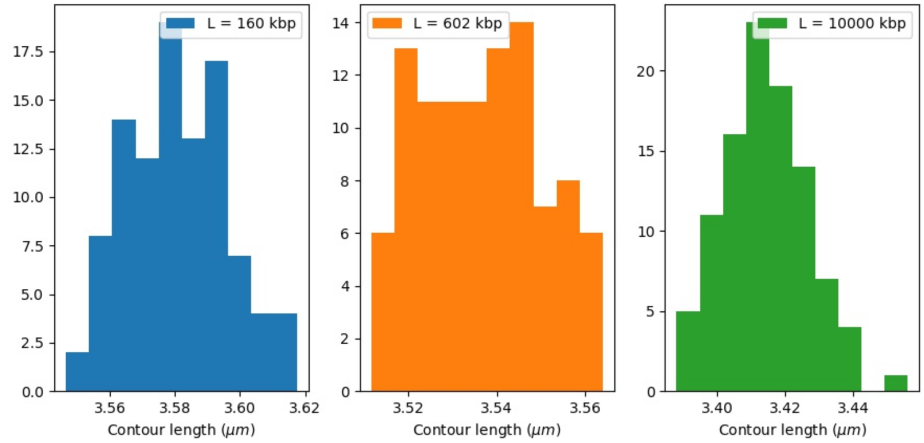


**Fig. S 9.** Distribution of contour lengths for a domain of size $L_d = 160$ kbp varying total polymer length (i) $L = L_d = 160$ kbp (ii) $L = 602$ kbp and (iii) $L = 10,000$ kbp. In all simulations volumic fraction $\phi = 0.055$ was kept constant.
